# SingloCell: an open and accessible web platform for multimodal single-cell data exploration

**DOI:** 10.64898/2026.11.23.767908

**Authors:** Alexis Allot, Hayden Wu, Elia Afanasiev, Moein Yaqubi, Sherilyn Junelle Recinto, Cameron Oram, Souren Farimani, Yue Zhang, Finn Creeggan, Josh Gertsvolf, Jiannis Ragoussis, Corina Nagy, Jun Ding, Sali Farhan, Jean-Francois Poulin, Jo Anne Stratton

**Affiliations:** Department of Neurology and Neurosurgery, Montreal Neurological Institute-Hospital, McGill University, Montreal, Quebec, H3A 2B4, Canada; McGill Genome Centre, Victor Phillip Dahdaleh Institute of Genomic Medicine, McGill University, Montreal, Quebec, H3A 0G1, Canada; Department of Psychiatry, McGill University, Montreal, Quebec, H3A 1A1, Canada; McGill Group for Suicide Studies, Douglas Mental Health University Institute, McGill University, Montreal, Quebec, H4H 1R3, Canada; School of Computer Science, McGill University, Montreal, Quebec H3A 2A7, Canada; Meakins-Christie Laboratories, Translational Research in Respiratory Diseases Program, Research Institute of the McGill University Health Centre, Montreal, Quebec H4A 3J1, Canada; Mila - Quebec Artificial Intelligence Institute, Montreal, Quebec H2S 3H1, Canada

## Abstract

Single-cell research is transforming biology and medicine, offering unprecedented insight into health and disease. Yet, the true potential of single-cell data can only be realized when datasets are open, easy to interact with, and accessible to all. We introduce SingloCell (https://singlocell.openscience.mcgill.ca/), an intuitive multimodal single-cell web platform built on three principles: accessibility, interactivity and traceability. SingloCell provides free browser-based access to single-cell datasets and supports user submission of suspension-based transcriptomics, proteomics, chromatin accessibility (ATAC-derived gene activity matrices), immune receptor (BCR, TCR) and multi-omics modalities, as well as spatial datasets. This enables researchers, clinicians, educators, and the broader community to explore high-dimensional datasets without programming expertise. Users can explore and analyse data in real time using tools for differential gene expression, functional enrichment, and pathway visualisation, while also examining pre-computed analyses such as pseudotime trajectories and cell-cell interactions. Datasets can be shared publicly or via unlisted links, and analytical workflows are documented through downloadable receipts.

**GRAPHICAL ABSTRACT:** 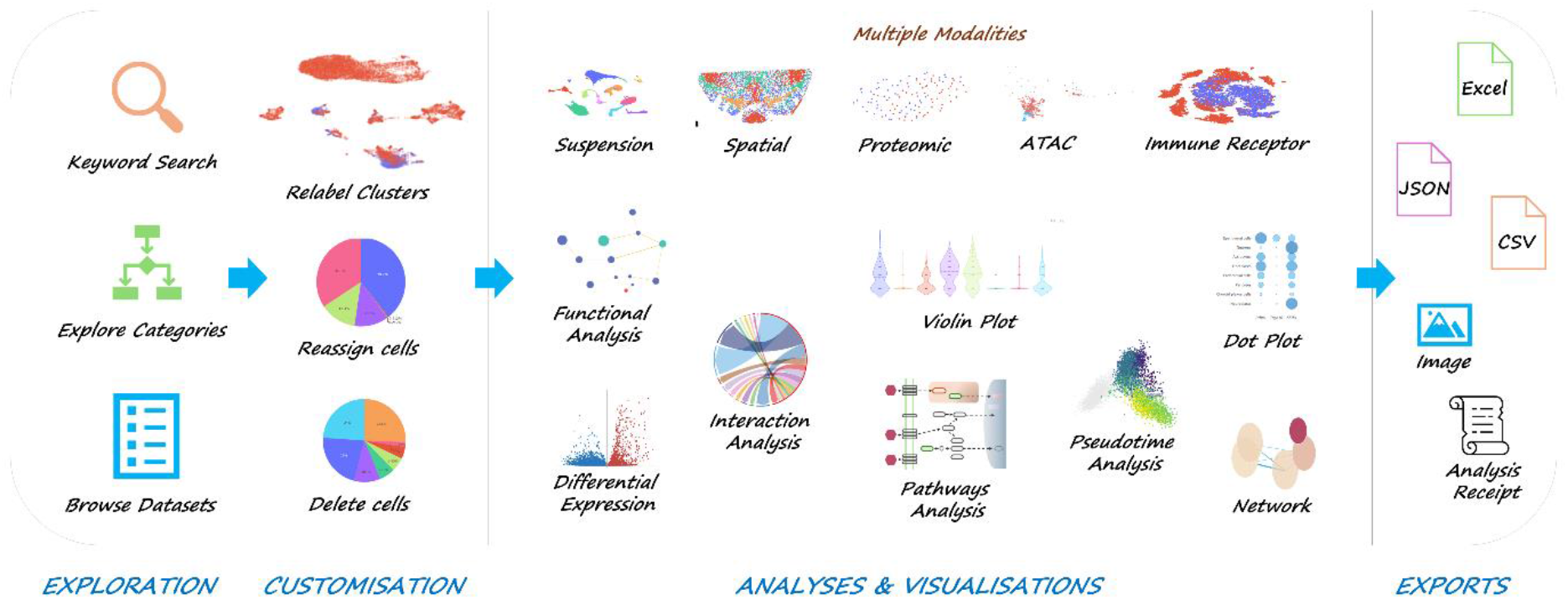

## INTRODUCTION

Single-cell research has rapidly transformed the life sciences, providing deep insights into cellular heterogeneity and disease mechanisms. It advances our understanding of diseases like cancer (Nam et al., 2021), neurological disorders (Groh et al., 2024), and immune responses (Qi et al., 2022), and is expected to shape the future of clinical diagnostics (Zhang et al., 2023) and personalized medicine (Ianevski et al., 2024; Tang et al., 2023). This rapid transformation is reflected by the increasing number of openly available online single-cell publications (PubMed)(Wilbrey-Clark et al., 2020) and datasets (GEO DataSets) (Clough et al., 2024).

In response to this increasing volume, numerous analysis platforms (Gondal et al., 2024), such as UCSC Cell Browser (Speir et al., 2021) and Trailmaker (Parse Biosciences, n.d.) have emerged. However, these resources do not always meet all open-science goals. Access is sometimes restricted, interfaces can be challenging to navigate, reusability and transparency can be limited, and dataset submission processes can be complicated or unavailable. This can limit collaborative discovery, broad accessibility, or reproducible analysis. A closer look at available web tools reveals a fundamental divide: display-oriented platforms which provide interactive views of datasets but limited interactivity and reanalysis (e.g., CZ CELLxGENE (CZI Cell Science Program et al., 2025)), and analysis-oriented platforms which offer advanced workflows, but often require technical expertise (e.g., Trailmaker (Parse Biosciences, n.d.)). These limitations become increasingly important as single-cell data becomes central to academic, industry and clinical applications. To address them, we believe that principles of accessibility, traceability and interactivity should be at the centre of online single-cell data analysis tools.

First, accessibility is key: platforms should provide free access to datasets of diverse modalities and present them in ways that are understandable to both specialists and non-specialists to improve communication and collaboration. Contributions should be straightforward, with platforms allowing users to submit new datasets through free, user-friendly workflows. Exploring available datasets should be simple and intuitive, with easy modifications, filtering and exporting. Second, traceability is critical to improve reproducibility. Settings used to generate analyses and visualisations should be available to users. The platform’s source code should also be open and publicly available. Finally, users should be able to go beyond static datasets. Platforms should provide interactive tools to explore, modify, and reanalyse existing datasets in real time, enabling users to easily repurpose them for new research projects.

To put these principles into practice, we created SingloCell (https://singlocell.openscience.mcgill.ca/), an open-source web platform for exploring, reusing and sharing curated single-cell datasets. Researchers can submit datasets either to accompany their publications through open, searchable links or to share through unlisted links, and no login is required to explore public datasets. SingloCell supports suspension-based transcriptomics, proteomics, chromatin accessibility (ATAC-derived gene activity matrices), immune receptor (BCR, TCR) and multi-omics modalities, as well as spatial datasets. We describe its architecture and feature below and compare it with existing platforms in Table 1.

**Table 1:** Popular single-cell analysis platforms. Platforms were evaluated against seven criteria grouped into three categories: accessibility, traceability and interactivity. These criteria were set to explore differences in sharing, exploring, analysing and documenting single-cell data. “Yes” means a criterion is fully met, “Partial” that it is met in part, and “No” that it is not met. Platform information accessed in November 2026.

| Criteria | **CZ CELLxGENE**<br>(CZI Cell Science Program et al., 2025) | **Trailmaker**<br>(Parse Biosciences, n.d.) | **UCSC Cell Browser**<br>(Speir et al., 2021) | **Single Cell Expression Atlas**<br>(George et al., 2024) | **Single Cell Portal**<br>(Tarhan et al., 2023) | **ABC Atlas**<br>(Yao et al., 2023) | **SingloCell** |
|---|---|---|---|---|---|---|---|
| **Accessibility** |  |  |  |  |  |  |  |
| Easy and free submission of diverse datasets | Partial<sup>a</sup> | No<sup>b</sup> | Partial<sup>a</sup> | Partial<sup>c</sup> | No<sup>b</sup> | No<sup>d</sup> | Yes |
| Open and easy dataset search | Yes | No<sup>e</sup> | Yes | Yes | Yes | Partial<sup>f</sup> | Yes |
| Pre-processing | Submitter | Website | Submitter | Submitter and website | Submitter | Website | Submitter |
| **Traceability** |  |  |  |  |  |  |  |
| Analysis Receipts | No | Partial<sup>g</sup> | No | No | No | No | Yes |
| Publicly available source code | Yes | Partial<sup>h</sup> | Yes | Yes | Yes | Partial<sup>i</sup> | Yes |
| **Interactivity** |  |  |  |  |  |  |  |
| Real time analysis | Partial<sup>j</sup> | Yes | No | No | Partial<sup>j</sup> | No | Yes |
| Adaptable and customisable | Yes | Yes | Yes | No | Partial<sup>k</sup> | No | Yes |

**Definitions:**
**Easy and free submission of *diverse* datasets:** Any user can submit datasets at no cost, without requiring an account or consortium affiliation. Platforms are scored Yes if they allow free, guided submissions with support for multiple data modalities. Platforms that do not accept submissions from external researchers or require an account or a fee are scored No. Platforms that rely on submissions to an external repository followed by a request for inclusion, or support only a limited range of data modalities or data types, are scored Partial.
**Open and easy dataset search:** Published datasets can be found without requiring an account. Platforms are scored Yes if search is available without login, Partial if search is limited to dataset titles, and No if login is required.
**Pre-processing:** “Submitter” means the dataset provider performs preprocessing and quality control before submission, and the platform applies no standard QC pipeline. “Website” means the platform itself has capacity for pre-processing of the data.
**Analysis receipt:** Analyses performed on the platform can be documented through an exportable record of the analysis parameters and software versions used. Platforms are scored Yes if an analysis receipt can be downloaded, Partial if analysis settings and steps are displayed but cannot be exported as a consolidated record, and No if no such information is provided.
**Publicly available source code:** Platforms are scored Yes if the complete source code is available, Partial if only part of the source code is publicly available, and No if no source code is publicly available.
**Real-time analysis:** Differential expression and functional enrichment analyses can be performed with results returned directly to the interface within a short time. Platforms are scored Yes if both analyses are supported, Partial if only one is supported, and No if neither is supported.
**Adaptable and customisable:** Users can modify the displayed dataset to personalise their own displays. Platforms are scored Yes if users can modify cell populations (e.g., by subsetting or regrouping), Partial if they can only annotate or label cells, and No if no modifications are supported.
**Notes:**
<sup>a</sup> Free submission is accepted after review by the platform team but largely restricted to RNA-seq modalities.
<sup>b</sup> A user account is required: Trailmaker requires an email sign-up (free for academic users and Parse Biosciences customers), and Single Cell Portal requires a Google account.
<sup>c</sup> Datasets must first be deposited in an EMBL-EBI archive, after which their inclusion in the Single Cell Expression Atlas is determined by the Atlas team. Only the RNA-seq modality is officially supported.
<sup>d</sup> Datasets are provided by the Allen Institute, and users cannot submit their own datasets.
<sup>e</sup> An account is required to access the dataset repository.
<sup>f</sup> Datasets can be searched by title only.
<sup>g</sup> Some analysis settings are documented, but a complete record of the analysis steps and parameters is not provided.
<sup>h</sup> Trailmaker is based on the open-source Cellenics platform (MIT licence), but it is unclear whether the released code matches the code used by the website.
<sup>i</sup> The web application’s code is not released. The available data-access code is under a licence that restricts commercial use and is therefore not open source.
<sup>j</sup> Only Differential Gene Expression is available.
<sup>k</sup> Only cell annotation and labelling is available.

## MATERIALS AND METHODS

### Architecture

The SingloCell website consists of two main components (Figure 1): an Angular-based frontend (Figure 1B) and a Django-based backend (Figure 1C). The frontend handles dataset visualisation, user interactions, and dataset management, providing an interface for exploring and modifying single-cell data. The backend is responsible for loading datasets and performing real-time analyses operating as a JSON-based REST API, meaning all endpoints return data in JSON format. This design ensures compatibility with alternative user-facing platforms, such as scripts, external applications, or third-party integrations, allowing broader accessibility and flexibility for advanced users.

**Figure 1:**
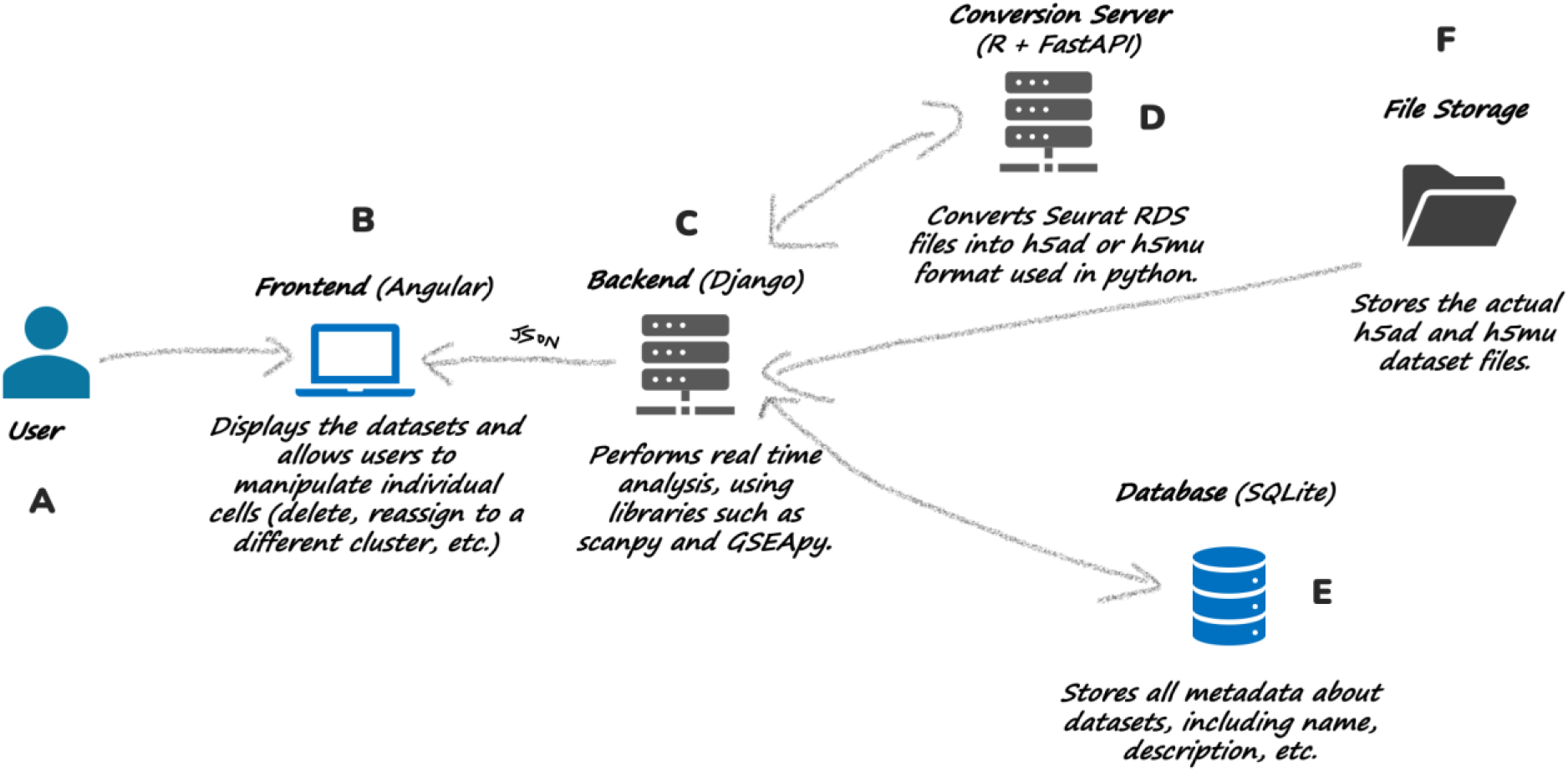
Server architecture. The SingloCell platform is composed of several components: the user (A) interacts via a web interface built with Angular (B), which displays datasets and allows manipulation of single-cell data. The frontend communicates with the backend server (C), which handles real-time analysis using libraries such as Scanpy and GSEApy. A conversion server (D) processes Seurat RDS files, converting them to h5ad or h5mu format required by Python-based tools. Metadata for each dataset is stored in a SQLite database (E), while the datasets themselves are stored as h5ad or h5mu files in file storage (F).

To enable efficient dataset searching, the frontend uses Fuse.js, which provides fast and flexible fuzzy searching. For data visualisation, we primarily use Plotly.js, which powers UMAP, dot plots, bar plots, violin plots, network graphs, volcano plots, heatmaps, and pathway visualisations. Plotly.js leverages WebGL, making it well-suited for rendering large datasets, such as displaying thousands of cells on UMAP plots. Additional visualisations, such as the chord diagram, are rendered directly using D3.js. When users modify datasets, such as renaming clusters, deleting cells, reassigning cells, or keeping only selected cells, the changes only affect the JSON representation maintained on the frontend. To perform real-time analysis, the frontend sends a list of selected cell IDs to the backend, ensuring efficient processing without modifying the original dataset files. Pathway visualisations are created by overlaying gene markers on static KEGG images, with gene positions extracted from KEGG’s XML files.

The backend is built using the Django framework and the Django REST Framework (DRF) to handle API requests. Dataset metadata is stored in an SQLite database, while the single-cell datasets themselves are stored on disk in h5ad or h5mu format. When a user requests to display a dataset, the backend loads the corresponding h5ad or h5mu file from disk using the anndata library and returns a JSON representation of the UMAP plot, generated with Plotly. The backend supports several real-time analyses, enabling interactive exploration of single-cell datasets. To compute DGE, after receiving a list of cell IDs from the frontend (representing target and background groups), the backend extracts the corresponding subset of the expression matrix. If the number of selected cells in either the target or background group exceeds a predefined threshold of 1000 cells, cells are subsampled using a cell-type-stratified uniform sampling strategy to optimize computational performance and enable real-time analysis while preserving representation of different selected cell populations. Genes expressed in fewer than 3 cells in either the target or background groups are removed. DGE analysis is then performed using Scanpy’s rank_genes_groups function with the Wilcoxon rank-sum test. Differential expression is run on the normalised expression values stored in the submitted object, and p-values are adjusted using the Benjamini-Hochberg procedure (Scanpy default). Genes are classified as significant based on an adjusted p-value below 0.05 and an absolute log2 fold change above 0.25. This does not account for patient/sample replicates and comparisons are intended as exploratory. Results are returned in JSON format for display on the frontend.

Functional enrichment analysis is performed on significantly differentially expressed genes using the Enrichr service through the GSEApy Python package. A list of enriched terms is returned to the frontend, along with a network of the top 10 terms constructed using the NetworkX library. Edge width represents the number of genes that overlap between two terms (nodes) while node size reflects the fraction of input genes overlapping with a term.

To overlay gene expression changes onto KEGG pathways, the backend stores pathway data in JSON format. This process includes retrieving pathway XML files from the KEGG API for human and mouse species, extracting pathway background images and encoding them in Base64, and storing gene coordinates relative to each pathway image. When the frontend requests a pathway visualisation with specific upregulated and downregulated genes, the backend retrieves the corresponding pre-computed JSON file and highlights the relevant genes.

SingloCell also supports the exploration of pre-computed cell–cell interactions, and pseudotime trajectories. Pre-computed cell–cell interaction data is added to the h5ad object as a pandas dataframe. When a user requests interactions between specific source and target cell types, the backend retrieves the relevant subset of data and returns it for visualisation on the frontend. Pre-computed pseudotime information is also stored in the dataset file by the dataset submitter. When a user makes a selection on the plot, the best matching pseudotime data is retrieved and displayed.

### Dataset Integration Pipeline and Deployment

All single-cell datasets submitted to SingloCell must be in Seurat RDS format or h5ad format. Once the file has been uploaded to the server, it is sent to an internal conversion service, running as a Docker container, which uses an in-house script to convert it to h5ad or h5mu format, if needed. If the conversion is successful, the h5ad or h5mu file is then automatically imported into the development server to be visualised.

Code updates are managed through an automated deployment pipeline (Figure 2). When a new update is pushed to the GitHub repository (Figure 2G), a TeamCity (Figure 2F) worker retrieves the changes, builds a Docker image, and deploys it on the server (Figure 2E). Both the frontend and backend have separate development and production branches, which are deployed as development and production websites, respectively. The development website is only accessible within the internal network, allowing new updates and datasets to be reviewed before they are publicly released. Once changes are validated, they are pushed to the production website, which is accessible to all users.

**Figure 2.**
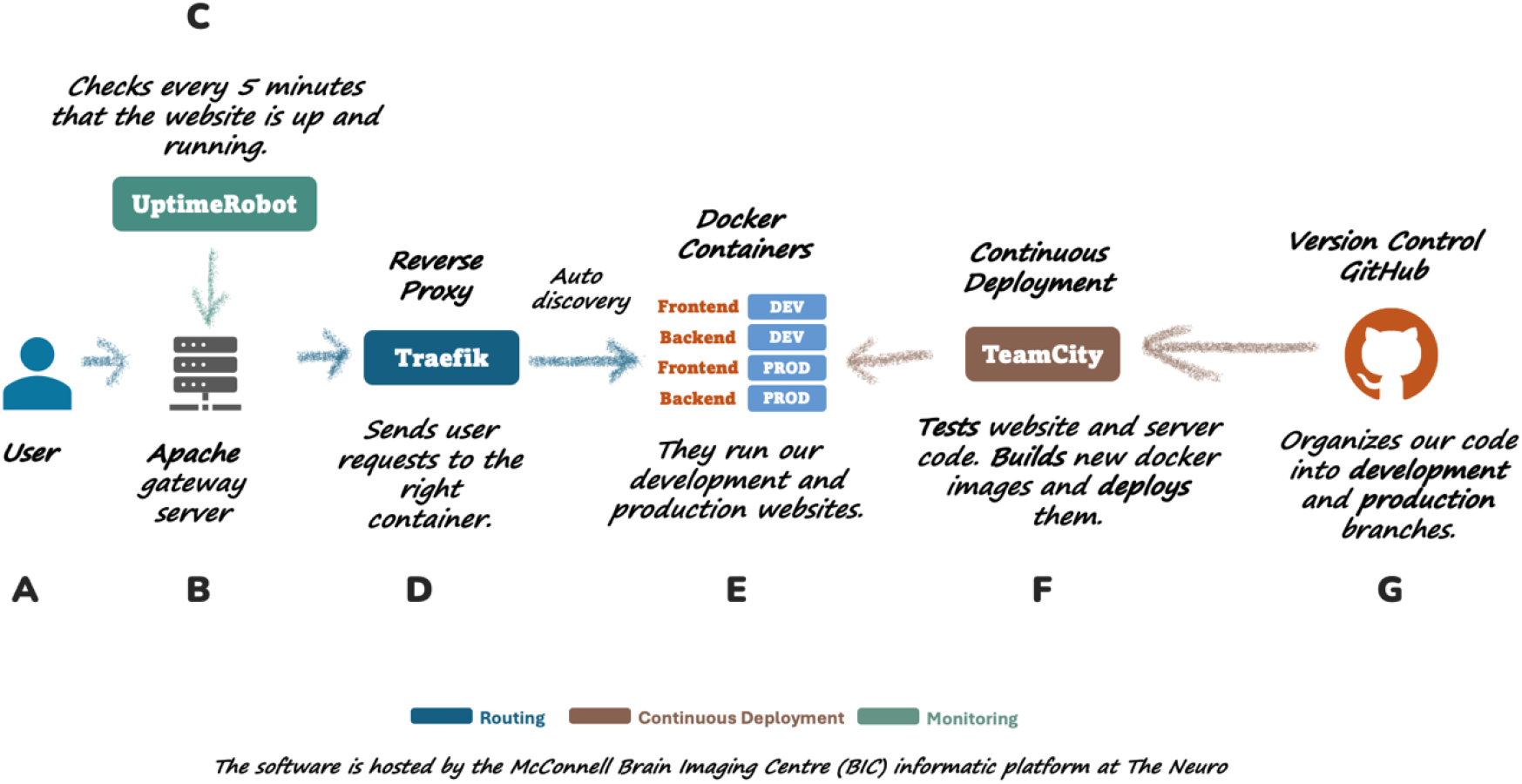
Overview of the SingloCell deployment architecture. User requests (A) are routed through the Apache gateway server (B) which is monitored by UptimeRobot (C) to ensure uptime. Traefik (D) acts as a reverse proxy, directing requests to the appropriate Docker containers (E), which run development and production versions of the frontend and backend. Continuous deployment is managed by TeamCity (F), which builds and deploys new Docker images from code stored and versioned on GitHub (G).

When a user (Figure 2A) accesses the website, their request is first routed through the gateway server (Figure 2B) and then passed to the Traefik reverse proxy (Figure 2D). Traefik directs the request to the appropriate Docker container based on the URL, ensuring access to the correct service. To ensure all websites are available and to receive alerts if any service becomes unresponsive, we use UptimeRobot (Figure 2C) as a monitoring solution.

### Code Availability

Source code is publicly available via GitHub (frontend is available at https://github.com/stratton-lab/cellar3-web-OPEN-SOURCE; backend is available at https://github.com/stratton-lab/cellar3-api-OPEN-SOURCE). Its architecture, data processing pipeline, and deployment are disclosed in this manuscript. The platform does not require paid software licences, is open source (MIT license), and its analytical methods rely on freely available Python packages described in the Materials and Methods section. Together, this provides an open and free foundation for adapting or independently deploying SingloCell.

## RESULTS

### Interactive Dataset Exploration and Customisation

SingloCell provides users with three flexible approaches to explore deposited datasets, catering to both broad exploration and specific searches.

The first approach relies on a curated category tree (Figure 3), which organises datasets into hierarchical classifications such as "*Suspension → Genomics → Baseline State"*, supporting exploratory browsing. This structure eases navigation, allowing users to locate datasets. For users who may not know exactly what they are looking for, the category tree makes it easy to browse and discover new datasets. The second approach is a search engine that enables precise dataset discovery. Users can enter queries that include dataset names, descriptions (containing terms such as “injury” or “immune”), species (such as “human” or “mouse”), tissues (like “brain” or “liver”), specific keywords (authors’ names, publication title, etc.), or experimental technologies (for example, “MERFISH”). Finally, the third approach allows users to filter datasets using a combination of attributes (e.g. Modality, State, Species, Tissue, etc.).

**Figure 3:**
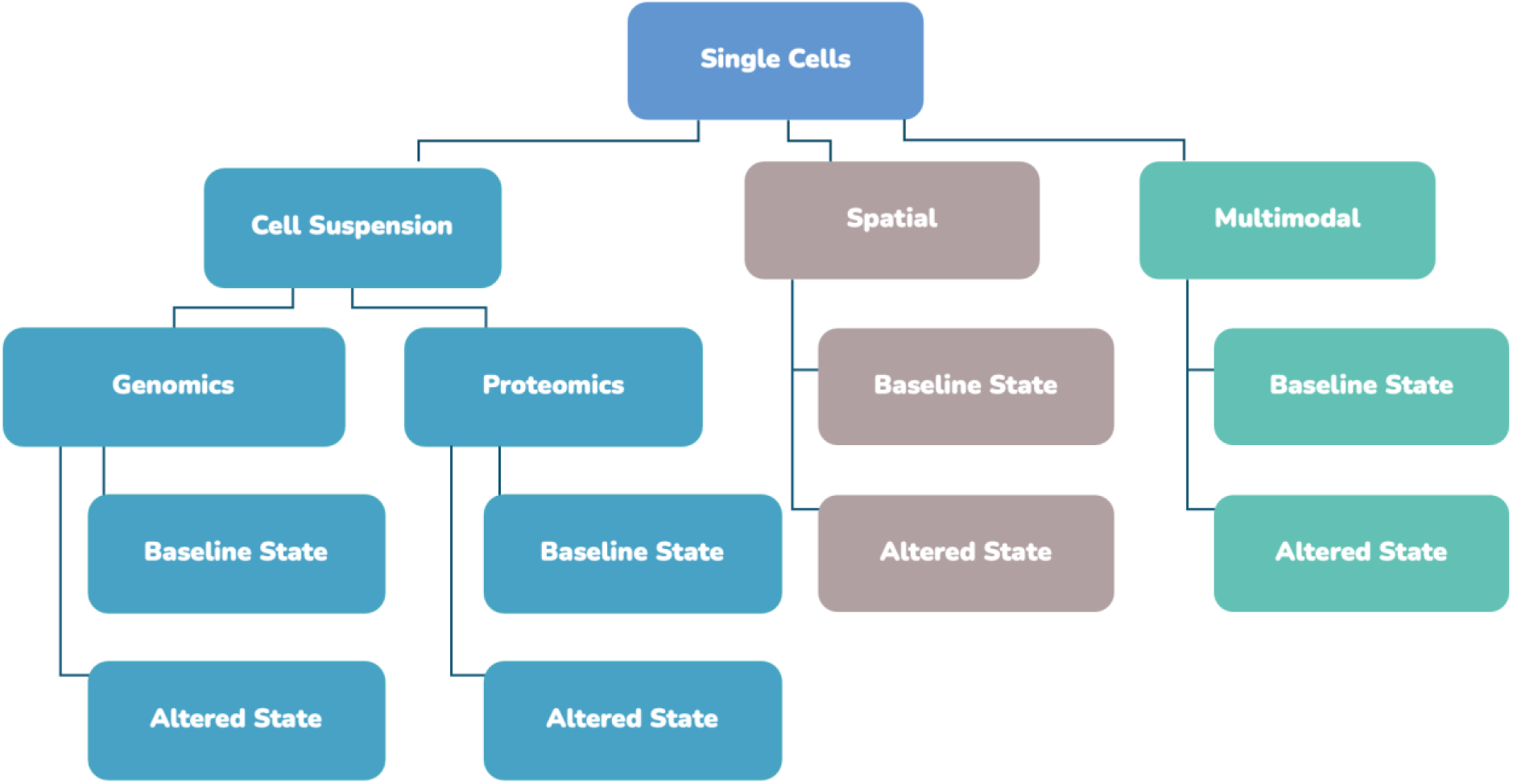
Current Categories Exploration Tree. SingloCell provides users with the ability to explore available datasets through nested categorizations.

After selecting a dataset, users are taken to a detailed dataset page (Figure 4A) that provides essential metadata, including the dataset’s name, description, tissue, organism, and cell count, as well as links to related publications and GEO or source data locations. The main feature of this page is an interactive plot displaying all the cells in the dataset. By default, cells are visualised using a UMAP embedding, while spatial datasets can be displayed according to their tissue coordinates. For multimodal datasets, the main interactive UMAP plot is accompanied by smaller previews of the other modalities, which automatically update to reflect cells removed from the main plot.

**Figure 4:**
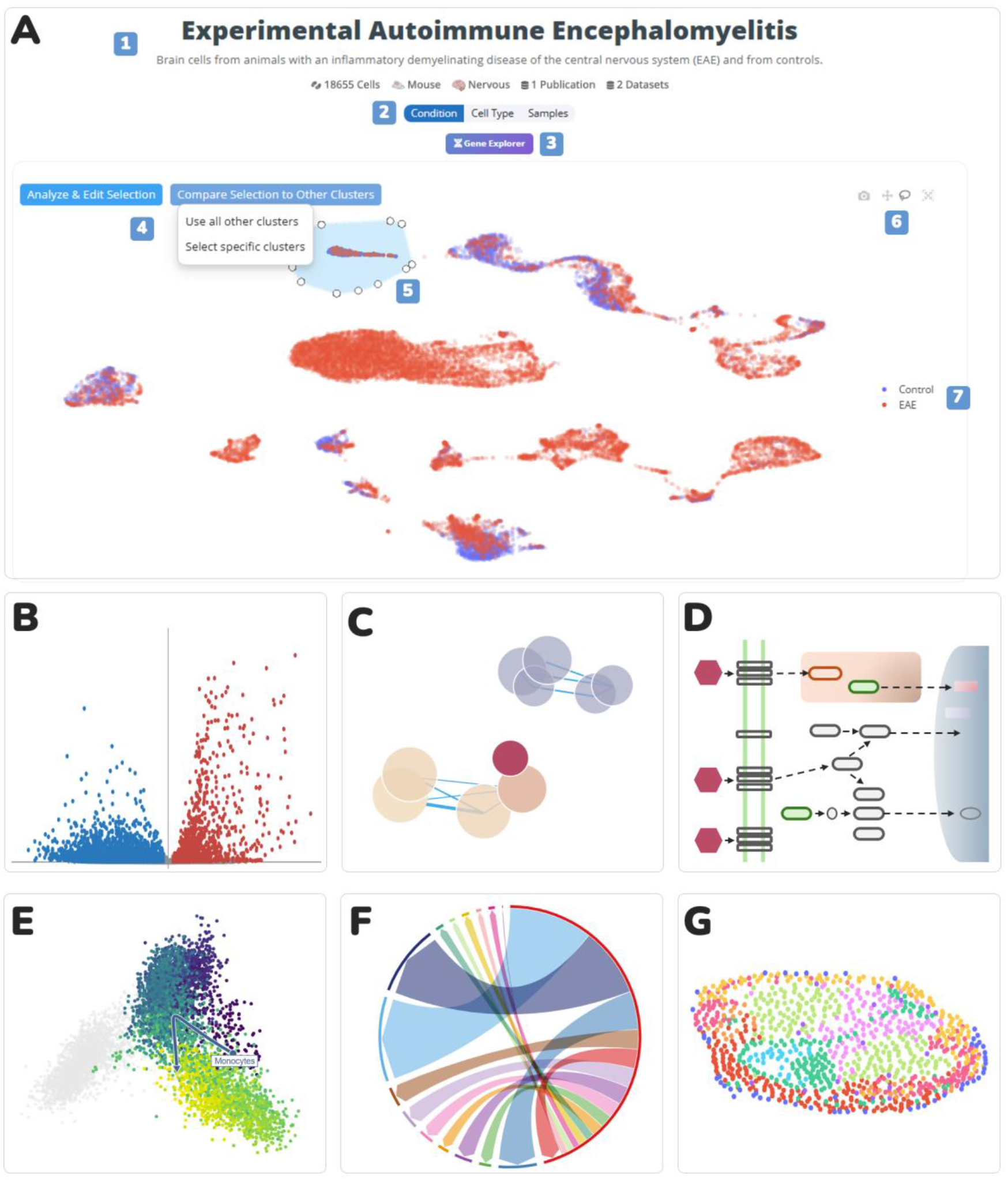
Interactive dataset page in SingloCell. Users can review dataset metadata (A1), group and colour cells by attributes such as condition or cell type (A2), launch the Feature Explorer for expression analysis (A3), run analyses and compare groups (A4), select cells directly on the main plot (A5), configure plot settings (A6), and customise cluster legends (A7). The interactive UMAP plot enables flexible selection, visualisation, and customisation of single-cell data. Available analyses include Differential Expression (B), Functional Enrichment (C), KEGG Pathway visualisation (D), Pseudotime trajectories (E), Cell-Cell interactions (F) and Spatial View (G).

The visualisation is fully interactive, allowing users to zoom, pan, and focus on specific regions of interest. Cells can be grouped and colour coded by attributes such as cell type, experimental condition, or sample origin, enabling researchers to quickly identify patterns and gain meaningful insights into cellular populations. For multimodal datasets, dedicated tabs allow users to switch between modalities, such as “RNA” or “Gene Activity” (ATAC).

SingloCell maintains a synchronised cell viewer across modalities, ensuring that interactive modifications are consistently reflected throughout the dataset. For example, cells deleted or retained in one modality are correspondingly updated across all other modalities. SingloCell also provides a "Feature Explorer" tool which allows users to investigate and compare the expression or gene activity levels of specific genes or proteins across the entire dataset. By entering gene or protein names into the search dialog, users can visualise their expression or gene activity patterns through violin, dot, and UMAP plots. This provides a detailed view of gene and protein levels across individual cells and modalities, enabling researchers to uncover meaningful biological insights.

SingloCell’s customisation features substantially enhance dataset reusability. Users can modify datasets by excluding irrelevant cells, merging samples, reassigning cells to new clusters, or keeping only selected cells. For example, cell types unrelated to a user’s research focus can be removed, simplifying the analysis. To encourage exploration and experimentation, the platform includes an "Undo" function, allowing users to revert changes.

Users can run analyses or compare selected groups with other cell populations in the dataset by selecting cells directly on the main plot. The plot can then be configured using action buttons in the top-right corner, allowing users to switch between drag and selection modes, recentre the view, or export the plot as a PNG image. Clusters can also be renamed, allowing users to tailor datasets to their specific needs. Further analyses are as shown in Figure 4B-G.

### Real Time Analysis

Differential gene expression, gene activity or protein expression is a common method for identifying targets that show significant differences in expression or gene activity between groups of cells and is central to many single-cell studies. SingloCell makes this process interactive by allowing users to perform comparisons between user-defined selected cells between samples and conditions, between a single group and the rest of the dataset, or among clusters within a selection. Results are displayed as an interactive volcano plot, highlighting significant features. From this plot, users can further explore patterns using violin, dot, and UMAP plots. Results can be downloaded in commonly used formats (CSV, Excel, JSON or PNG Image) and further explored in offline workflows. In addition to lists of significantly regulated features, excel format exports also include metadata describing cell IDs used in the analysis. For multi-omics datasets, significant regulation of all modalities is exported within this excel format.

Building on this, functional enrichment analysis in SingloCell identifies enriched biological categories linked to differentially expressed features between selected groups of cells. Users can examine results in categories such as Function and Pathways, Biological Process, Cellular Component, and Molecular Function. The results are visualised through bar plots or networks (Figure 4C), supporting the interpretation of complex relationships. The bar plot allows users to hover over diverse categories to view the number of upregulated and downregulated genes, while the network representation connects enriched categories based on overlapping genes.

For pathway-specific insights, SingloCell displays KEGG diagrams (Figure 4D) that highlight connections between significant features, offering a detailed view of their roles within larger biological processes.

Extending beyond expression and functional relationships, SingloCell allows users to study trajectories through pre-computed pseudotime data. Once a group of cells is selected, a pseudotime plot highlights their position along an inferred trajectory (Figure 4E). The visualisation includes arrows and labels to indicate the trajectory direction and origin, which are based on the predominant cell type within the selection. Users can further investigate expression patterns in the same embedding, with cells colour-coded to reflect the expression levels of selected features.

To complement trajectory analysis, SingloCell also provides tools to study pre-computed cell-cell interactions. After selecting cells of a specific type, users can generate a chord diagram that illustrates interactions between the predominant cell type in the selection and other cell types in the dataset (Figure 4F). This feature allows filtering by interaction type, such as "Secreted Signalling," enabling researchers to focus on specific cellular communication processes. Selecting a specific interaction between two cell types generates a heatmap that details ligand-receptor pairs, along with their associated probabilities. From the heatmap, users can explore individual interactions further by viewing supporting evidence via links to relevant publications or by visualising the ligand and receptor genes on a KEGG pathway diagram.

In addition to displaying cell suspension datasets, SingloCell also supports spatial data (Figure 4G), allowing users to explore individual cells in relation to their location within a tissue. Spatial datasets can be explored in two ways: in the regular integrated mode, where users view all cells together as a global overview, or in spatial mode, where users can select specific tissue slices to display and examine the physical arrangement of cells within the tissue context. Each spatial embedding includes an overview image highlighting the position of cells in the broader tissue section, helping users explore cellular states within the tissue architecture.

### Transparency and Traceability

SingloCell allows users to download an analysis receipt in JSON format. The receipt documents the dataset analysed, the Python version, and the analytical steps performed, including relevant libraries and parameters. For example, a DGE receipt records the subsampling settings, the number of cells before and after subsampling, the Scanpy version, and the statistical methods and thresholds used. The receipt is intended as a readable record of the analysis rather than an executable script. In addition to the receipt, one can export the DGE Excel, where the cell IDs that were used for a given analysis are listed.

As a test case, to reproduce findings purely from exported receipt and cell IDs, we ran dataset RNA_Ms_HeartLA_ALDO_2025 (30,193 cells in total). Within the dataset, we subset the endocardial cell cluster using the lasso tool and compared aldosterone-treated cells (n = 3,805) with control cells (n = 3,963). Each group exceeded the 1,000-cell limit, so SingloCell subsampled each to 1,000 cells. The subsampled cell IDs are listed in the exported Excel file and define the result, as repeating the comparison on the platform may select different cells. Offline, we re-ran the analysis with the Python and Scanpy versions recorded in the receipt (3.11.5 and 1.9.6), the parameters recorded in the receipt and described in Materials and Methods, and the cell IDs recorded in the Excel file. The results matched the platform for all 12,271 genes tested (maximum difference in log2 fold change below 10⁻¹⁵), including all 219 significant genes.

### Dataset Submissions

Submitting a new dataset to SingloCell is a guided process that begins with a pre-submission inquiry form (21 questions, requiring 15 minutes for a new user), where users provide key information about themselves and their dataset.

After the admin team reviews the pre-submission inquiry, the user receives an email with a link to an upload form, allowing them to submit their dataset as an RDS Seurat file or anndata h5ad file. Users can also upload a preview image for their dataset (300 × 300 pixels in size) which will be displayed (or an image will be generated by the admin team based on dataset description). Users submitting datasets are responsible for performing appropriate preprocessing, quality control, normalisation, and cell type annotation steps prior to submission: SingloCell does not enforce a universal QC pipeline or fixed filtering criteria for the submitted datasets. To ensure a consistent experience across devices and internet connections, submitted RDS files are limited to 2 GB in size (∼250K cells with ∼20K genes). Therefore, any additional assays and layers (e.g., count data) that are not needed for display, should be removed by the submitter prior to submission to improve upload reliability and reduce unnecessary data transfer.

For datasets containing chromatin accessibility data, submitters must provide a gene-by-cell gene activity matrix. As gene activity scores depend on the preprocessing pipeline and genomic regions considered (e.g., gene bodies, promoters or distal enhancers), submitters are encouraged to document their methods in the dataset description and use informative modality names.

Once uploaded, the file is automatically converted to h5ad (pre-2026) or h5mu (post-2026) format using SingloCell’s conversion service and deployed to a development environment for initial assessment. The admin team conducts a thorough review, and the dataset is then deployed to SingloCell’s production website in unlisted mode, where access is restricted to the submitter and any user they choose to share the access link with. Through an iterative process, the submitter can confirm that the data reproduces key findings from the original RDS Seurat file or anndata h5ad file that was submitted. The submitter can keep the data in private mode or, at any point, they can approve the dataset for public release at which point it becomes accessible to all users browsing or searching the website.

### Case Study: Exploration of Early Immune Dysregulation in PINK1 KO Infected Mouse

Growing evidence suggests that gastrointestinal (GI) tract pathology may contribute to the initiation and progression of Parkinson’s disease (PD), particularly through interactions between environmental triggers and genetic susceptibility. Pathogenic variants in PINK1, an established cause of early-onset PD, have been associated with enhanced proinflammatory profiles and altered immune responses, suggesting that immune dysregulation may emerge outside the central nervous system during early disease processes.

Using SingloCell, researchers can interactively explore single-cell transcriptomic data from PINK1 knockout (KO) mice following infection to investigate potential changes in immune cell populations and their transcriptional responses (Dataset ID: RNA_Ms_Gut_Immune_PD_2024). Users can visualise cellular heterogeneity through interactive UMAP plots and compare cell populations across experimental conditions and time points. For example, specific myeloid or T-cell populations can be isolated to examine differences in gene expression, identify candidate inflammatory pathways, and explore how these responses vary across experimental groups. These comparisons provide an accessible starting point for investigating how PINK1 deficiency may influence peripheral immune responses following an inflammatory challenge.

To investigate these responses further, SingloCell enables users to explore pre-computed pseudotime trajectories, revealing potential transcriptional transitions within selected immune populations. Genes associated with these inferred trajectories can be visualised to identify candidate intermediate cell states and pathways associated with immune activation. In parallel, pre-computed cell–cell interaction analyses allow users to examine predicted ligand–receptor interactions between immune populations, generating hypotheses about alterations in intercellular communication. Together, these tools illustrate how SingloCell can support initial data exploration of focused biological questions, facilitating the identification of candidate immune pathways and cellular interactions for further investigation in PD.

## DISCUSSION

Single-cell studies generate large and complex datasets, yet published results are often presented through static figures that offer limited opportunities for further exploration. Researchers seeking to explore these datasets further may need to download large files and use specialized software, requiring advanced computational expertise. SingloCell aims to overcome these barriers by providing a free and accessible web platform for interactive exploration of single-cell datasets.

Compared with existing single-cell platforms (Table 1), SingloCell places particular emphasis on accessibility, traceability, and interactivity, combining straightforward dataset submission, interactive analysis, and documentation of analytical workflows to facilitate the reuse of published data. Rather than serving as a comprehensive data repository, SingloCell allows individual researchers to upload curated datasets as interactive resources, for example in parallel with their publications, enabling readers to explore single-cell data beyond the analyses presented in the original study. It also does not provide a comprehensive single-cell analysis pipeline, from raw data to visualisation, as offered by platforms such as Trailmaker. Rather, SingloCell relies on datasets that have already undergone preprocessing, quality control, annotation, and precomputed analyses. This approach preserves the analytical decisions of the original study, allowing dataset submitters to ensure that the data presented on SingloCell reflect their published findings. However, this means that data quality rests with the submitters. We also prioritise rapid, real-time exploration through subsampling to enable a streamlined experience. Users requiring more comprehensive analyses can export the metadata of selected cells and perform further, more granular analyses locally. For donor-aware testing, users can export the selected cell IDs and run pseudobulk analysis on the original counts from the source data. Finally, although multimodal datasets (e.g. RNA and ATAC-derived gene activity measured in the same cell) are supported, SingloCell does not provide comprehensive integration of independently generated datasets. These design choices reflect a deliberate trade-off favouring interactive exploration over exhaustive analysis, positioning SingloCell as a complement to more comprehensive analytical workflows.

Future developments will focus on improving scalability to support larger datasets and reduce the need for subsampling, expanding analytical capabilities across modalities, and refining the user interface based on community feedback. We also plan to support additional submission formats to encourage contributions from a broader range of researchers. By combining accessible data exploration with reproducible analytical workflows, SingloCell aims to support open science and enable researchers from diverse backgrounds to investigate new biological and medical questions using existing single-cell datasets.

## ACKNOWLEDGEMENTS

AI tools were used during the development of the SingloCell web server and preparation of the manuscript to assist with code writing, text drafting and editing, and the creation of visual assets. All scientific interpretations, analyses, and final decisions were made by the authors.

## AUTHOR CONTRIBUTIONS

A.A. developed the SingloCell website. A.A. and H.W. tested all components, proposed new features, provided feedback on usability, and managed dataset submissions, processing, and communication with dataset owners. E.A. helped develop the dataset conversion script. H.W., M.Y., S.J.R., C.O., So.F., Y.Z., F.C., and J.G. uploaded and curated datasets on the website. A.A., H.W., and J.A.S. drafted the manuscript. J.A.S. conceived and supervised the project. J.R., C.N., J.D., Sa.F., and J.F.P. provided project funding and intellectual contributions.

## DATA AVAILABILITY

SingloCell is free and open to all users and there is no login requirement. It is available at https://singlocell.openscience.mcgill.ca/. The code for the website frontend is available at https://github.com/stratton-lab/cellar3-web-OPEN-SOURCE The code for the website backend is available at https://github.com/stratton-lab/cellar3-api-OPEN-SOURCE

